# Adjuvanted mucosal vaccination enhances protection and prevents influenza virus transmission in the guinea pig model

**DOI:** 10.64898/2026.08.03.742639

**Authors:** Vivian Yan, Seok-Chan Park, Matthew J. Wiest, Gabriel Laghlali, Julianne N. d’Acunzo, Chinghan Chung, Jorge Levican, Farah El-Ayache, Pamela T. Wong, Michael Schotsaert

**Affiliations:** Department of Microbiology, Icahn School of Medicine at Mount Sinai, New York, NY, 10029, USA; Global Health and Emerging Pathogens Institute, Icahn School of Medicine at Mount Sinai, New York, NY, 10029, USA; Graduate School of Biomedical Sciences, Icahn School of Medicine at Mount Sinai, New York, New York, USA; Department of Internal Medicine, University of Michigan Medical School, Ann Arbor, Michigan, USA; Michigan Nanotechnology Institute for Medicine and Biological Sciences, University of Michigan Medical School, Ann Arbor, MI, USA; Department of Pharmaceutical Sciences, Ghent University, Ghent, Belgium; Department of Biomedical Engineering, University of Michigan, Ann Arbor, Michigan, USA; Icahn Genomics Institute, Icahn School of Medicine at Mount Sinai, New York, New York, USA; Marc and Jennifer Lipschultz Institute for Precision Immunology, Icahn School of Medicine at Mount Sinai, New York, New York, USA; Department of Immunology and Immunotherapy, Icahn School of Medicine at Mount Sinai, New York, New York, USA

## Abstract

Influenza virus infects the respiratory mucosa, highlighting the importance of mucosal immunity for early protection and transmission control. Here, we evaluated whether intranasal (IN) vaccination with recombinant trimeric hemagglutinin protein from A/Michigan/45/2015 (triHA) formulated with a combined mucosal adjuvant, nanoemulsion plus IVT, an RNA-based RIG-I agonist (NE/IVT), could protect guinea pigs against heterologous A/Netherlands/602/2009 challenge and reduce viral transmission. To compare mucosal and parenteral immunization, IN triHA/NE/IVT was benchmarked against IN triHA alone, IM triHA/AddaVax (IM triHA/Advx), and standard IM quadrivalent inactivated influenza vaccine (QIV). We also tested whether IN triHA/NE/IVT could boost IM QIV- primed immunity and included animals previously infected with A/Michigan/45/2015 to model pre-existing infection-induced immunity. Transmission was assessed by co-housing naïve sentinels with vaccinated, challenged donors. IN triHA/NE/IVT induced systemic humoral responses comparable to IM triHA/Advx while generating superior nasal mucosal IgA responses. Unexpectedly, IM triHA/AddaVax also induced detectable, albeit lower, mucosal IgG and IgA, contrasting with prior mouse data and highlighting species-specific differences. IN triHA/NE/IVT boosting after IM QIV enhanced serum IgG and mucosal IgA compared with QIV prime-boost alone and increased cross-neutralizing activity against antigenically distinct A/Victoria/4897/2022. Both IN triHA/NE/IVT and IN Michigan/15 prior-infection prevented detectable viral shedding after challenge, and naïve sentinels co-housed with IN triHA/NE/IVT- vaccinated donors remained seronegative. Together, these findings support NE/IVT as a potential mucosal platform capable of inducing robust systemic and mucosal immunity and boosting IM vaccine-primed responses.

## Introduction

Influenza virus primarily infects mucosal surfaces of the respiratory tract and continues to impose a major global health burden. Although infection- or vaccine-induced immunity can reduce disease severity, influenza viruses frequently evade pre-existing antibody responses through antigenic drift or shift, allowing breakthrough infection, viral replication, and onward transmission to susceptible hosts (1–5). In particular, systemic immunity induced by conventional intramuscular (IM) vaccination may be insufficient to fully control viral replication at mucosal surfaces, where initial infection and shedding occur. As a result, individuals with vaccine induced immunity may still support viral replication in the upper respiratory tract and contribute to transmission (1, 2, 5). Thus, protection from severe disease does not necessarily equate to prevention of mucosal virus replication or transmission, highlighting the need for vaccine strategies that strengthen immunity at the site of viral entry and shedding.

Inducing immunity directly at respiratory mucosal sites may therefore provide an important advantage over systemic immunity alone (1, 2, 5). Local mucosal immune responses can act early after exposure by limiting viral entry, reducing productive replication, and suppressing shedding of newly produced virions from the upper respiratory tract (6–9). Because respiratory viral shedding is a key driver of influenza transmission, vaccines capable of strengthening mucosal immunity may help reduce both individual disease burden and population-level spread. This is particularly relevant for antigenically drifted influenza variants, which can transmit even in previously vaccinated populations (10).

In this study, we evaluate a recombinant protein vaccine delivered intranasally (IN) with a rationally designed mucosal adjuvant, a nanoemulsion (NE) formulation possessing TLR-and NLRP3 agonistic properties combined with a retinoic acid-inducible gene I (RIG-I) agonist (11–13). The RIG-I agonist is a 5’ prime phosphorylated *in vitro* transcribed (IVT) copy back defective interfering RNA that is produced during Sendai virus infection and is a strong inducer of Type I interferon responses as described by *Patel et al.* (14). We hypothesize that, similar to natural infection, simultaneously targeting multiple innate immune pathways at mucosal sites will result in induction of better qualitative immune responses both in the periphery (serum) and the mucosa, resulting in blockage of direct infection and viral transmission. We have evaluated the use of the combination adjuvant NE/IVT in mice and hamsters using recombinant SARS-CoV-2 spike protein vaccines, in which robust induction of systemic as well as mucosal antibody and T cell responses were observed within the sera and respiratory tract, respectively (12, 13).

We conducted this study in the guinea pig influenza model, which enables evaluation of both vaccine-mediated protection in directly infected animals and transmission to naïve sentinel animals (15–18). Because guinea pigs are susceptible to unadapted human influenza viruses and support efficient respiratory droplet transmission, this model is well suited for assessing viral shedding and interhost spread. Using recombinant trimeric hemagglutinin (triHA) as the vaccine antigen, we evaluated whether NE/IVT-adjuvanted IN vaccination could enhance systemic and mucosal immunity, limit viral replication, and reduce transmission. We compared this approach with established parenteral strategies, including IM triHA with AddaVax and standard IM quadrivalent inactivated influenza vaccine (QIV), and assessed its potential as a mucosal booster after QIV priming. We also included animals with prior H1N1 infection to compare vaccine-induced and infection-induced protection. Together, our results show that NE/IVT-adjuvanted IN vaccination induced systemic humoral immunity comparable to parenteral vaccination while preferentially enhancing mucosal IgA responses. This was associated with reduced viral replication, attenuated lung pathology, and marked inhibition of interhost transmission. NE/IVT also effectively boosted IM QIV-primed responses, supporting its potential as a flexible platform for next-generation influenza vaccines.

## Results

### Rationally Designed Combined Mucosal Adjuvants Elicit Robust Systemic and Superior Local Immunity in the Guinea Pig Model

We first assessed the transmissibility of A/Michigan/45/2015 (Michigan/15), the homologous strain used to design the triHA vaccine antigen. Donor guinea pigs were IN inoculated with 1 × 10^4^ PFU of Michigan/15 and co-housed with naïve sentinels 8 h later. Although all donors shed infectious virus by 3 DPI and viral RNA was detected from 1–5 DPI, no infectious virus was recovered from sentinels, and only one sentinel showed transient viral RNA detection at 3 DPI (Supplement 1A). Because Michigan/15 transmitted inefficiently in this setting, A/Netherlands/602/2009 (NL/09) was selected for the main challenge and transmission studies.

To investigate the efficacy of the NE/IVT combination adjuvant, guinea pigs were assigned to experimental groups (n = 3–4 per group) and immunized via either the IN or IM route (Fig. 1A, B). A control group (IN-PBS) received IN phosphate-buffered saline at three-week intervals. For systemic comparison, one group received recombinant triHA protein derived from A/Michigan/45/2015 formulated with the MF59-equivalent IM adjuvant AddaVax (IM triHA/Advx). Mucosal vaccination groups included unadjuvanted IN triHA and IN triHA/NE/IVT. In addition, two QIV-based groups were included, using a formulation containing an A/Victoria/4897/2022 (Victoria/22)-like H1N1 component: an IM QIV; IN triHA/NE/IVT group, in which IM QIV priming was followed by a heterologous IN boost with adjuvanted triHA, and an IM QIV prime-boost group. To simulate pre-existing immunity from natural infection, an IN Michigan/15 group was included, in which animals were IN infected with A/Michigan/45/2015 seven weeks before the final challenge. Four weeks after the second boost, or seven weeks after the initial Michigan/15 infection, all animals were IN challenged with 1×10^4^ PFU of A/Netherlands/602/2009 (H1N1) to evaluate the magnitude and quality of protection.

**Figure 1.**
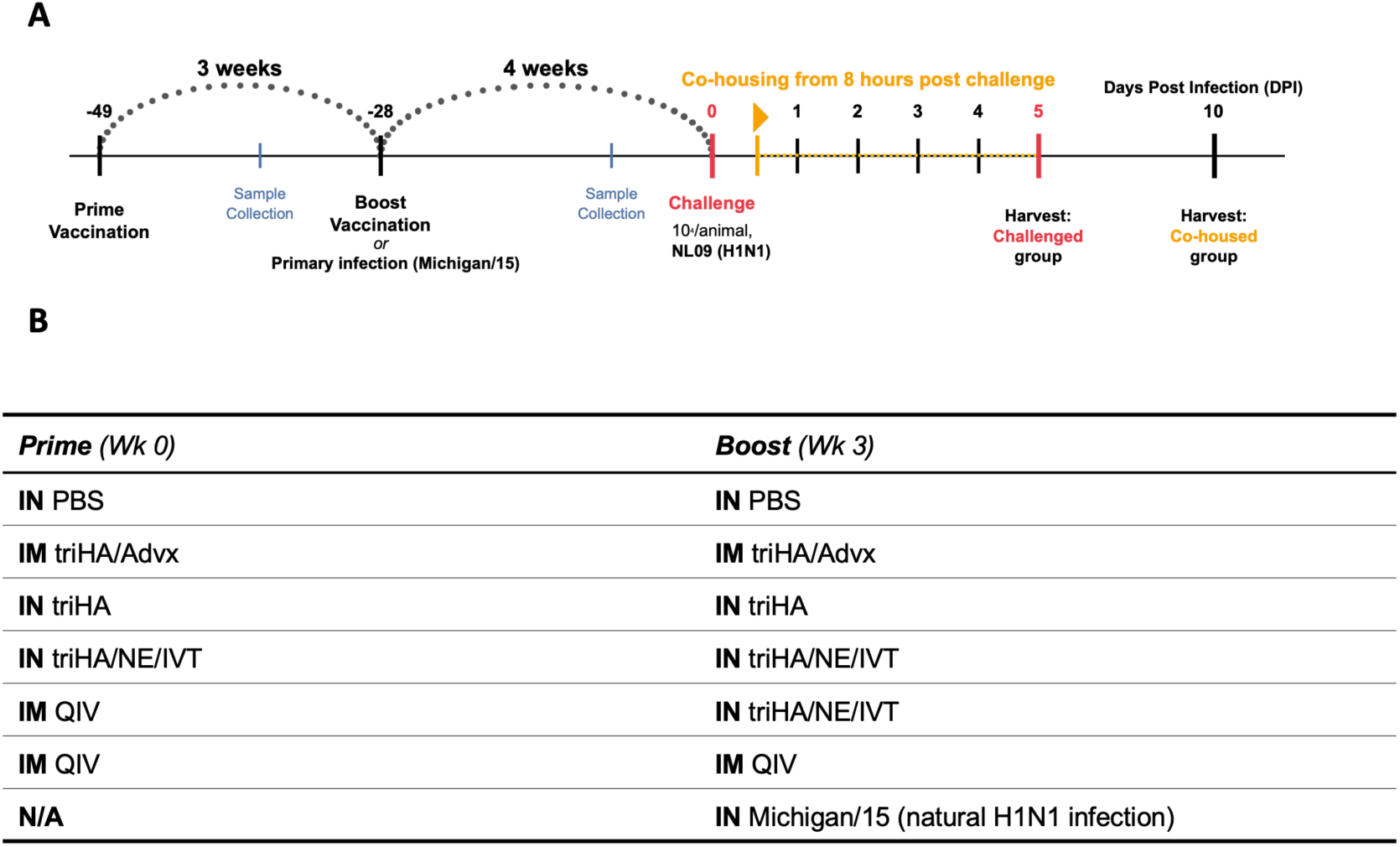
Experimental framework and transmission kinetics of A/Michigan/45/2015. (A) Vaccination and challenge study design. Guinea pigs were assigned to experimental groups and received prime and boost immunizations 21 days apart using the formulations summarized in (B). Twenty-eight days after boost immunization, animals were intranasally challenged with A/Netherlands/602/2009. To evaluate transmission, one naïve, unvaccinated sentinel animal was introduced into the cage of each challenged donor 8 h after infection and co-housed for 5 days. Nasal washes were collected longitudinally to assess viral shedding and transmission. (C) Summary of vaccination groups, routes, antigen formulations, and challenge conditions used in the study.

Two weeks after prime vaccination, 100% seroconversion was observed only in the IM triHA/Advx and IN triHA/NE/IVT groups, whereas unadjuvanted IN triHA failed to induce detectable responses and the QIV groups showed only limited seroconversion (Fig. 2A). After boost, IM triHA/Advx induced the highest serum IgG titers, followed by IN triHA/NE/IVT, showing that NE/IVT-adjuvanted IN vaccination can generate strong systemic humoral responses despite mucosal delivery (Fig. 2B). The IM QIV; IN triHA/NE/IVT group also showed higher serum IgG than the IM QIV prime-boost group, indicating that mucosal boosting enhanced QIV-primed responses. Serum IgA was undetectable after prime but became detectable after boost in 100% of the IM triHA/Advx group and in 50% of the IN triHA, IN triHA/NE/IVT, and IM QIV; IN triHA/NE/IVT groups (Fig. 2C, D).

**Figure 2.**
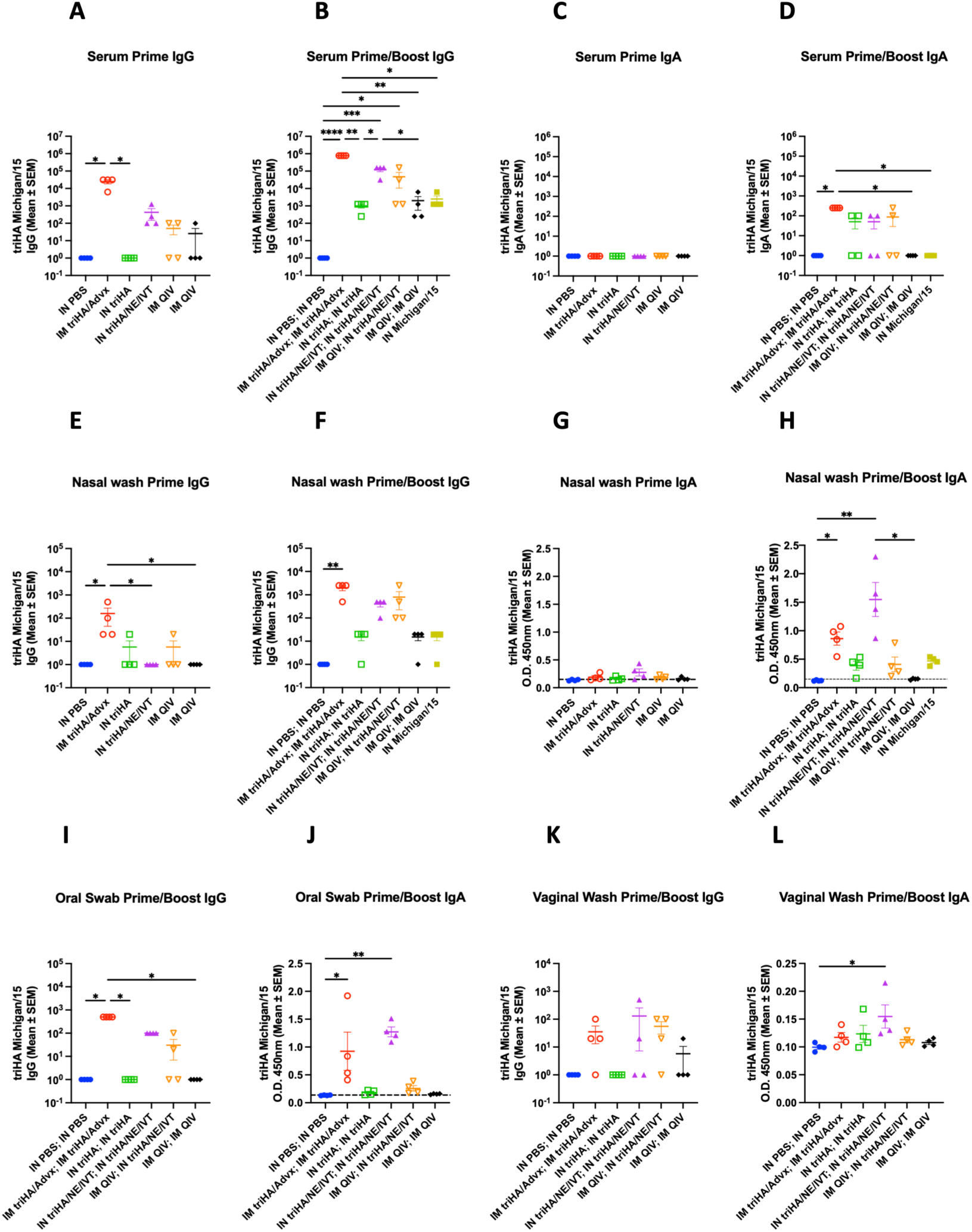
Antibody responses in serum and at mucosal sites post vaccinati on. Animals were swabbed for mucosal secretions and bled for serum collection. Sera and mucosal samples were analyzed by ELISA for (B–E) HA-specific serum IgG (B, C) and IgA (D, E) responses following prime and boost immunization, (F–I) HA-specific IgG (F, G) and IgA (H, I) in nasal washes obtained after prime and boost, and (J–M) HA-specific IgG (J, K) and IgA (L, M) titers in oral swabs and vaginal wa shes collected after the boost immunization. Data are shown for individual animals with group means ± SEM/95% CI . The dashed line indicates the limit of detection of the assay. Statistical significance was determined using a Kruskal–Wallis test followed by Dunn’s multiple-comparison test; *, P < 0.05; **, P < 0.01; ***, P < 0.001; ****, P < 0.0001.

Analysis of nasal washes revealed distinct mucosal antibody profiles (Fig. 2E–H). Nasal IgG largely mirrored serum responses, with the highest titers in the IM triHA/Advx group, followed by IM QIV; IN triHA/NE/IVT and IN triHA/NE/IVT (Fig. 2E, F). In contrast, nasal IgA was most strongly induced by IN triHA/NE/IVT after boost and was significantly higher than in the IN PBS and IM QIV groups (Fig. 2H). Importantly, IM QIV priming followed by IN triHA/NE/IVT boosting also markedly increased nasal IgA compared with IM QIV prime-boost alone, supporting the use of mucosal boosting to overcome the limited mucosal immunity induced by parenteral vaccination. Notably, IM triHA/Advx also induced modest nasal IgA, in contrast to prior mouse studies.

To assess distal mucosal responses, we also examined oral swabs and vaginal washes (Fig. 2I–L). IM triHA/Advx induced the strongest IgG responses in oral samples (Fig. 2I), whereas IN triHA/NE/IVT induced the most robust IgA responses, including significantly increased IgA levels in both oral swabs and vaginal washes compared with the IN PBS group (Fig. 2 J, L). Together, these findings show that NE/IVT-adjuvanted IN vaccination elicits strong systemic humoral immunity while preferentially enhancing local and distal mucosal IgA responses in guinea pigs.

### Mucosal Adjuvants Expand Neutralizing Antibody Breadth and Mitigate Antigen Mismatch in Influenza Vaccination

To assess antibody breadth and functional activity, post-boost sera were analyzed by microneutralization against NL/09, Michigan/15, and the antigenically drifted Victoria/22 (Fig. 3A–C). Consistent with the serum IgG data, the IM triHA/Advx group generated the highest neutralizing titers against NL/09 and Michigan/15, followed by the IN triHA/NE/IVT and IM QIV; IN triHA/NE/IVT groups. The IM QIV and IN Michigan/15 groups showed similar neutralizing activity against these two strains. In contrast, unadjuvanted IN triHA induced little or no neutralizing activity, indicating that NE/IVT improved both the magnitude and functional quality of the antibody response. Neutralization was markedly reduced against Victoria/22 in most groups (Fig. 3C), with MNT titers decreasing by 20- to 80-fold relative to NL/09 and Michigan/15. This reduction was less pronounced in the QIV-based groups, which included a Victoria/22-like H1N1 component. Notably, the IM QIV; IN triHA/NE/IVT group showed stronger neutralizing activity against Victoria/22 than the matched IM QIV prime-boost group, suggesting that heterologous IN boosting with triHA/NE/IVT can enhance cross-neutralizing breadth against antigenically drifted strains.

**Figure 3.**
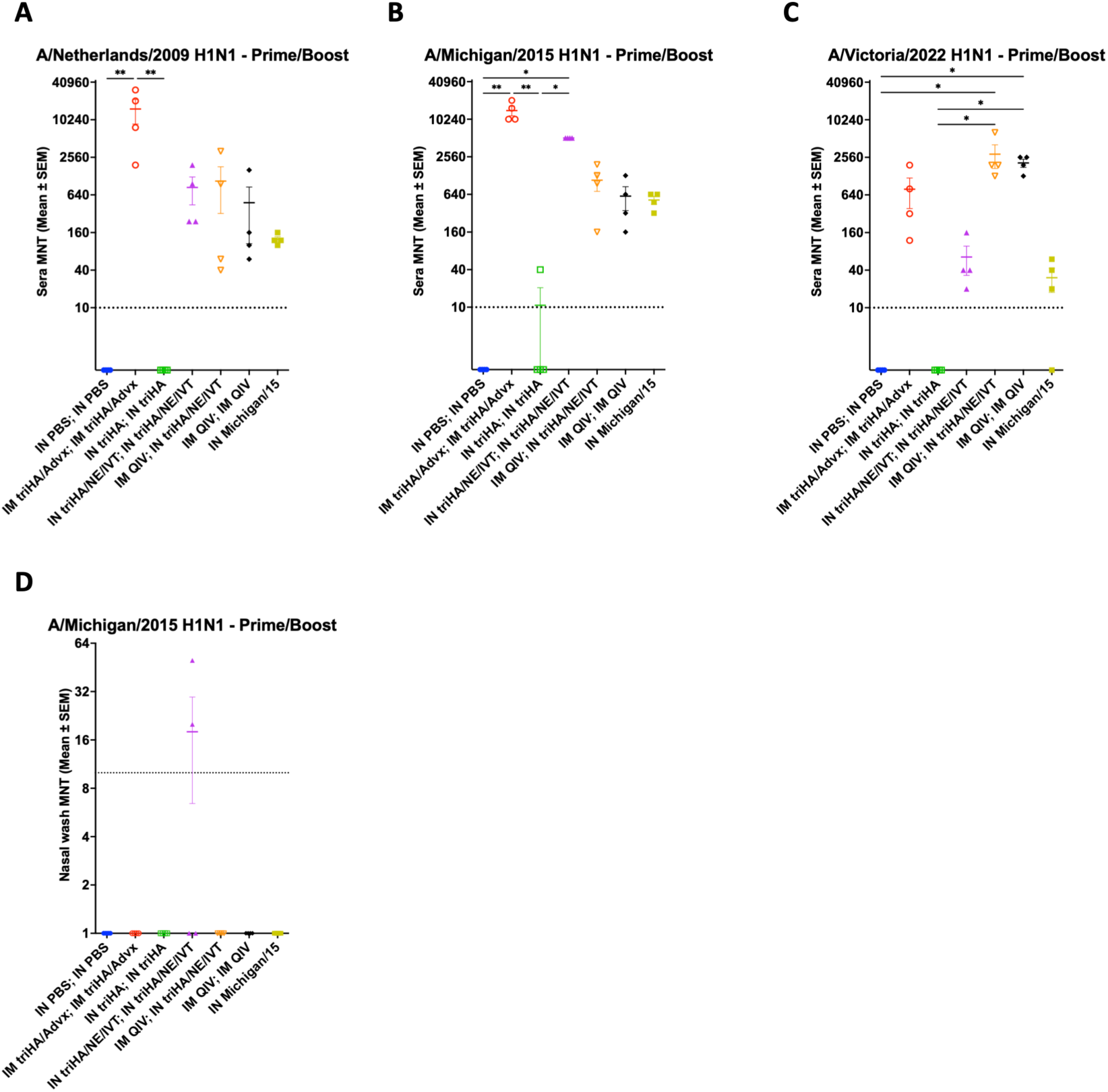
Serum microneutralization titers against homologous and heterologous H1N1 influenza viruses. Sera collected from vaccinated guinea pigs after boost vaccination were assessed in a microneutralization assay against (A) the challenge strain A/Netherlands/602/2009 (H1N1), (B) A/Michigan/45/2015 (H1N1) which antigenically matches the recombinant triHA vaccine protein, and (C) A/Victoria/4897/2022 (H1N1) which is antigenically matching the QIV H1N1 strain. (D) Nasal wash samples collected after boost vaccination were assessed by microneutralization assay against A/Michigan/45/2015 (H1N1). Reciprocal microneutralization titers from individual animals in each vaccination group are shown, with geometric mean titers ± standard error of the mean. The dashed line indicates the limit of detection of the assay. Statistical significance was determined using a Kruskal–Wallis nonparametric one-way ANOVA followed by Dunn’s multiple-comparison test with multiplicity-adjusted P values; *, P < 0.05; **, P < 0.01;

Microneutralization assays were also performed on nasal wash samples to assess upper respiratory tract neutralizing activity. No detectable neutralizing activity against NL/09 or Victoria/22 was observed in nasal washes from any group (data not shown). In contrast, nasal washes from two animals in the IN triHA/NE/IVT group showed detectable neutralizing activity against Michigan/15 (Fig. 3D).

### Adjuvanted Mucosal Immunization Significantly Reduces Histopathologic Lung Lesions Despite the Subclinical Nature of IAV Infection in Guinea Pigs

To evaluate the protective efficacy of the vaccine candidates against viral pathogenicity and transmission, vaccinated guinea pigs were IN challenged with NL/09 four weeks after booster immunization. Eight hours after challenge, naïve sentinel animals were introduced into the index cages for direct-contact transmission studies. Clinical signs were monitored in challenged donor animals from 1 to 5 days post-infection (DPI). Consistent with the guinea pig model as a largely subclinical model of influenza infection, no animals developed severe morbidity (Fig. 4A). Modest weight loss was observed in the IM triHA/Advx group, which reached approximately 5.3% at 2 DPI and partially recovered by 5 DPI. The unadjuvanted IN triHA group also showed a mild reduction in body weight, whereas the remaining vaccinated groups and naïve sentinels remained clinically unremarkable throughout the study period (Fig. 4A, B).

**Figure 4.**
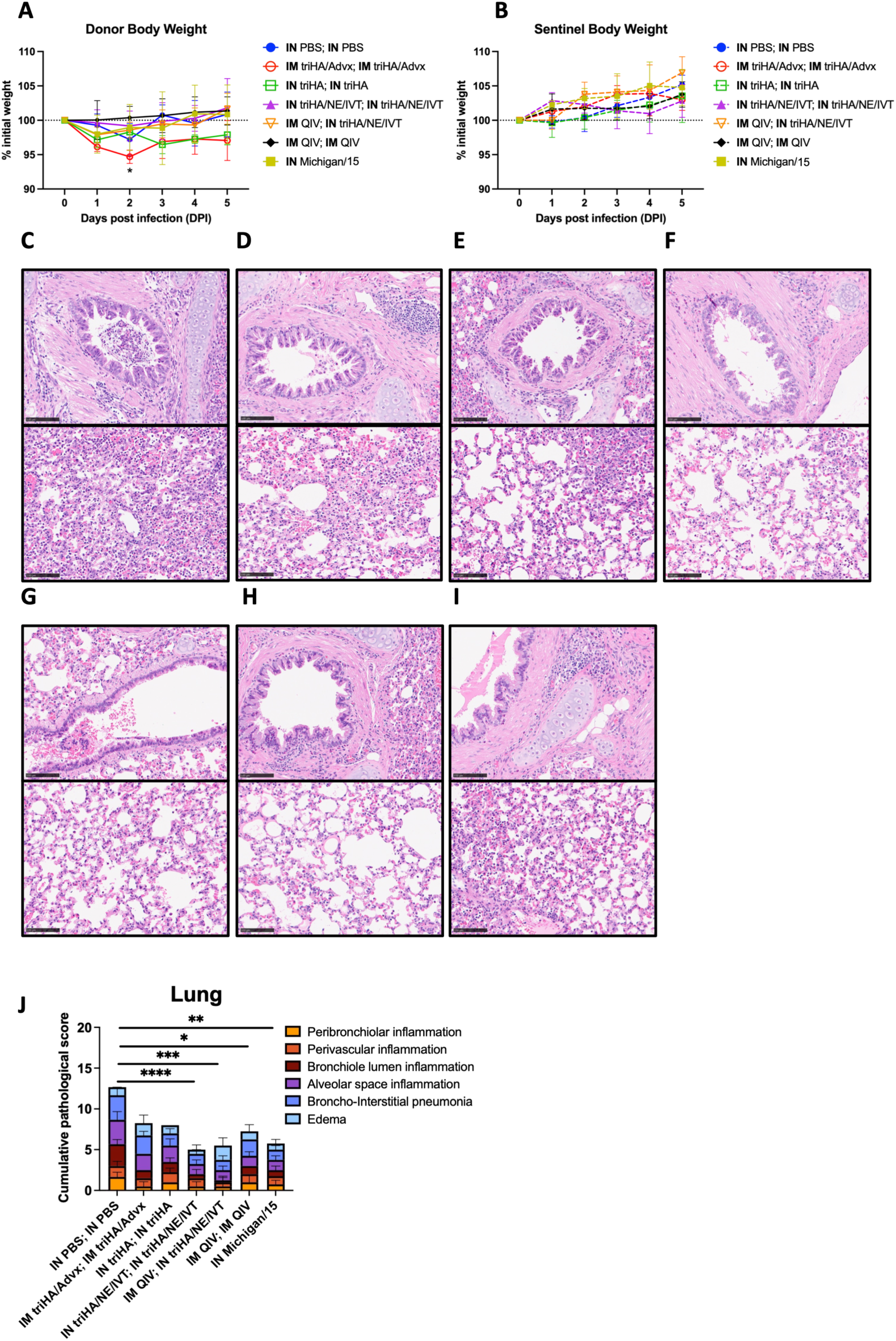
Body weight loss, lung pathology, and lower respiratory tract antibody responses after heterologous challenge. (A) Body weight changes in vaccinated donor guinea pigs and naïve sentinel (B) guinea pigs. Animals vaccinated as indicated were intranasally challenged with 10⁴ PFU of A/Netherlands/602/2009 influenza virus; naïve sentinels were co-housed with each vaccinated and challenged donor beginning 8 h post-infection for 5 days. Body weight was monitored daily and is expressed as the mean percentage of the day 0 weight ± SEM (n = 4 per group). Body weight at 1 to 5 days post-infection was compared between donors and sentinels using two-tailed Welch’s t tests with Holm-Šidák correction for five comparisons; days 1 to 5 remained significant after correction (multiplicity-adjusted P values of 0.0019, 0.0019, 0.0048, 0.0037, and 0.0113, respectively; donors < sentinels at each time point). (C–I) Representative hematoxylin-and-eosin–stained lung sections from donor guinea pigs at 5 days post- infection. Shown are lungs from animals in the IN mock (PBS) group (C), IM triHA/Adda<u>V</u>vax group (D), IN triHA group (E), IN triHA/NE/IVT group (F), IM QIV followed by IN triHA/NE/IVT group (G), two-dose IM QIV group (G) and IN Michigan/15 infection group (I) and corresponding cumulative pathological scores (J) for the respective vaccination groups, based on scoring of peribronchiolar, perivascular, bronchiolar luminal, and alveolar inflammation consistent with broncho-interstitial pneumonia. Horizontal bars indicate group means ± SEM. Asterisks indicate P values as determined by the tests described above: *, P < 0.05; **, P < 0.01; ***, P < 0.001; ****, P < 0.0001.

To determine whether vaccination reduced respiratory tissue damage despite the limited overt clinical disease in this model, histopathological analysis was performed on lungs collected at 5 DPI (Fig. 4C–I). The IN PBS control group showed characteristic lesions, including mild peribronchiolar inflammation with mononuclear cell infiltration and mild-to-moderate broncho-interstitial pneumonia (Fig. 4C). In contrast, the IM triHA/Advx group showed improved histopathologic findings relative to IN PBS controls despite mild clinical weight loss (Fig. 4D). Quantitative analysis of cumulative histopathology scores showed that the IN triHA/NE/IVT, IM QIV; IN triHA/NE/IVT, IM QIV, and IN Michigan/15 groups all had significantly reduced lung pathology compared with non-vaccinated controls (Fig. 4J). Although mild peribronchiolar inflammatory cell infiltration was observed in some vaccinated animals, no significant differences were detected among the vaccinated groups (Fig. 4E–I). Together, these data show that vaccine-induced protection in the guinea pig model is more readily detected by histopathologic analysis than by clinical signs alone, and that both mucosal and systemic vaccination reduced virus-associated lung lesions after challenge.

### Superior Blockade of Viral Shedding and Interhost Transmission via Adjuvanted Mucosal Vaccination

Nasal washes were collected from vaccinated, NL/09-challenged donor animals and co-housed naïve sentinels to assess viral shedding and contact transmission. Infectious virus was quantified by plaque assay, and viral RNA was measured by qPCR. The IN PBS group showed sustained donor shedding from 1 to 5 DPI, and two of three sentinels became infected, confirming efficient transmission (Fig. 5A). The IM triHA/Advx group showed substantially reduced donor shedding, with infectious virus detected in only one of four donors, and no sentinel animals became plaque-positive, although one of four sentinels later seroconverted (Fig. 5B, H). In contrast, unadjuvanted IN triHA provided limited protection: all donors shed infectious virus, and all sentinels became infected (Fig. 5C, H).

**Figure 5.**
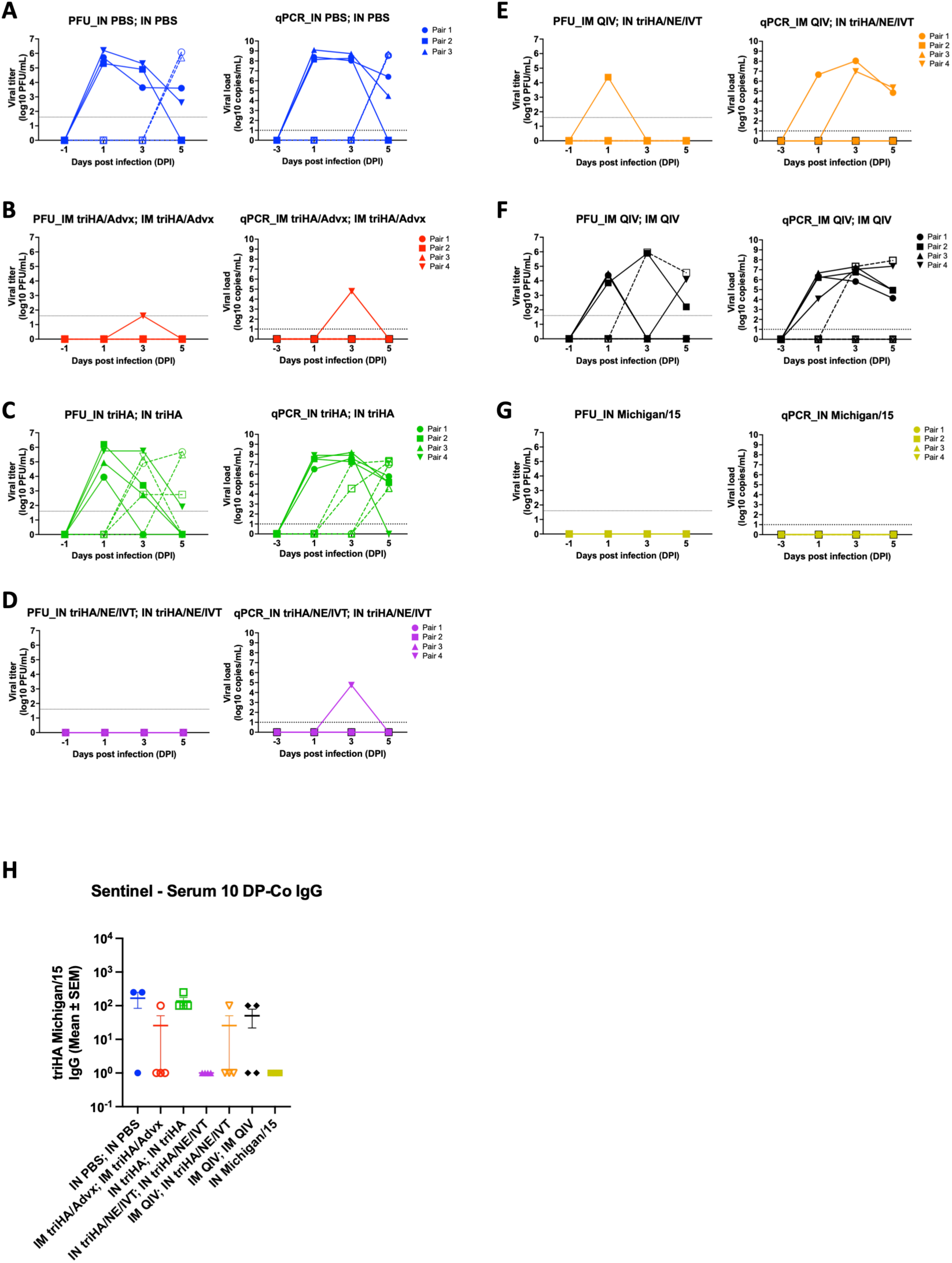
Intranasal vaccination with mucosal adjuvants reduces viral shedding and prevents transmission in a direct-contact guinea pig model. Vaccinated donor guinea pigs were challenged intranasally and co-housed with naïve sentinel animals to assess viral shedding and transmission. Nasal washes were collected from donor and sentinel animals at 1, 3, and 5 days post-infection (dpi). Infectious virus was quantified by plaque assay, and total viral RNA was measured by qPCR. Sera from sentinel animals were collected 10 days after the end of co-housing to assess seroconversion. (A–G) Viral shedding in donor and sentinel animals from the (A) IN PBS, (B) IM triHA/Advx, (C) IN triHA, (D) IN triHA/NE/IVT, (E) IM QIV; IN triHA/NE/IVT, (F) IM QIV, and (G) IN Michigan/15 groups. For each group, the left panel shows infectious viral titers measured by plaque assay, and the right panel shows viral RNA levels measured by qPCR. (H) Sentinel seroconversion measured 10 days after co-housing. Donor animals are shown as filled symbols with solid lines, and sentinel animals are shown as open symbols with dashed lines. Dashed horizontal lines indicate the assay limit of detection where applicable.

IN triHA/NE/IVT provided the strongest protection. No infectious virus was recovered from any donor or sentinel animal, and qPCR detected only a transient signal in one donor at 3 DPI, with no sentinel seroconversion, indicating complete blockade of detectable transmission (Fig. 5D, H). The IM QIV; IN triHA/NE/IVT group also showed strong protection, with infectious virus detected in only one donor at 1 DPI and no plaque-positive sentinels, although one sentinel seroconverted, indicating partial exposure despite marked transmission reduction (Fig. 5E, H). IM QIV alone provided only partial protection: all donors shed infectious virus, and one of four sentinels became plaque-positive, while two of four seroconverted (Fig. 5F, H). The IN Michigan/15 group also showed complete protection, with no detectable donor shedding, no infected sentinels, and no sentinel seroconversion (Fig. 5G, H). Together, these results show that IN triHA/NE/IVT most effectively reduced donor shedding and prevented transmission, whereas unadjuvanted IN triHA failed to protect and IM QIV alone provided only partial control.

### Addavax Adjuvant Mediates Robust Systemic and Respiratory IgA Responses Independent of Antigenic Context in Guinea Pigs

IM triHA/Advx elicited unexpectedly high sera and nasal wash IgA titers compared to IN PBS—a pattern unreported in mouse studies (19). This finding was notable because parenteral vaccination is generally a suboptimal inducer of mucosal immunity in mice and typically fails to generate detectable respiratory secretory IgA. Because this response was observed after vaccination with influenza HA, we next asked whether the effect was specific to the influenza antigen or reflected a broader feature of IM AddaVax adjuvantation in guinea pigs. To address this, we immunized guinea pigs with an unrelated antigen, trimeric SARS-CoV-2 Spike protein, with or without AddaVax (IM triS and IM triS/AddaVax, respectively). Two weeks post-boost, IM triS/AddaVax similarly induced elevated IgG and IgA in both sera and nasal washes (Fig. 6A-D), demonstrating that the response is platform-driven rather than antigen- specific. This suggests guinea pigs mount distinct mucosal and systemic antibody responses to IM/AddaVax vaccination, highlighting the necessity of multiple animal models to capture immunological diversity relevant to human vaccine design.

**Figure 6.**
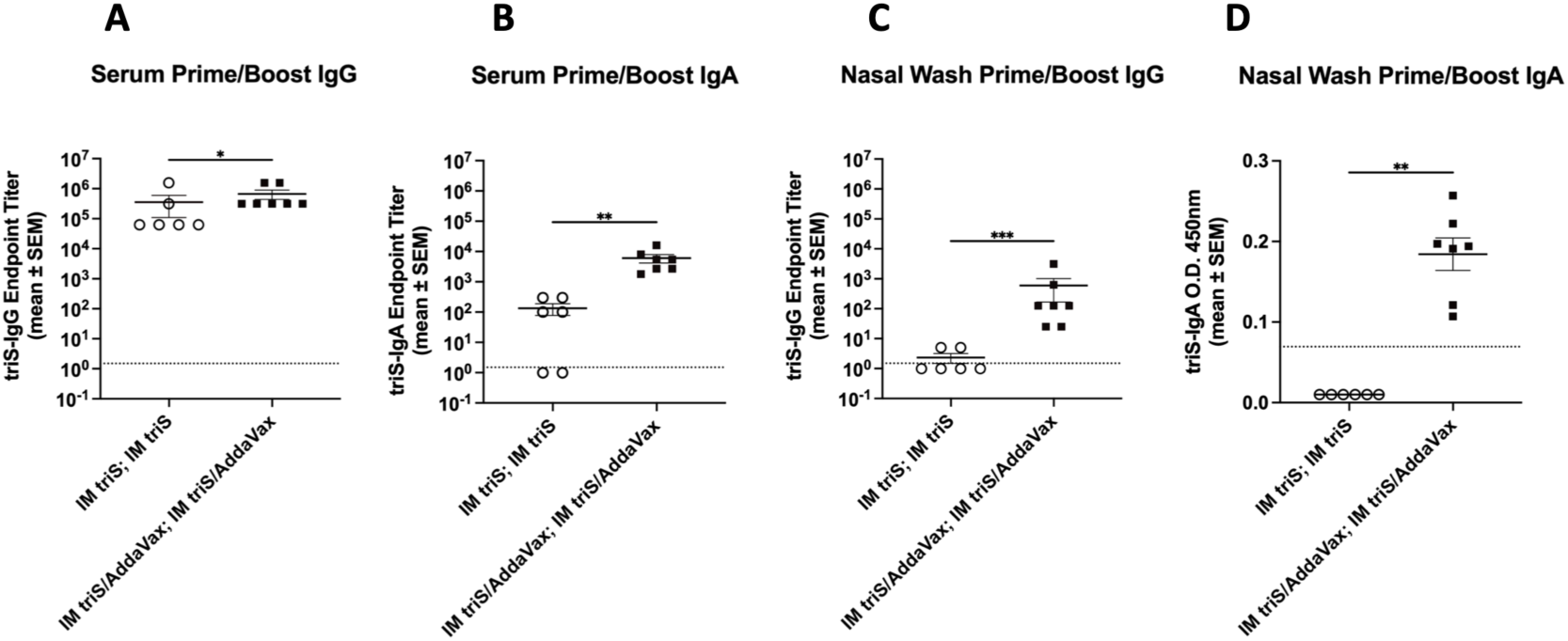
Intramuscular AddaVax-adjuvanted vaccination with an unrelated triSpike antigen induces systemic and mucosal antibody responses in guinea pigs. Serum and nasal wash samples were collected at the indicated time point after boost immunization to assess whether IM adjuvanted vaccination could promote mucosal antibody responses. TriSpike-specific antibody titers were measured by ELISA for (A) serum IgG, (B) serum IgA, (C) nasal wash IgG, and (D) nasal wash IgA. Each symbol represents an individual animal; circles indicate guinea pigs vaccinated IM with triSpike alone, and squares indicate guinea pigs vaccinated IM with triSpike formulated with AddaVax. Data are shown as mean ± SEM. Statistical significance was determined using a Mann–Whitney test. Asterisks indicate P values as follows: *, P < 0.05; **, P < 0.01; ***, P < 0.001; ****, P < 0.0001.

## Discussion

In this study, we evaluated an IN mucosal vaccination strategy using the rationally designed NE/IVT combination adjuvant in the guinea pig influenza model. This approach was compared with established IM regimens, including AddaVax-adjuvanted triHA and QIV, as well as prior infection. Previous murine studies showed that NE/IVT activates multiple innate immune pathways and induces robust systemic and mucosal humoral immunity (11, 13). Building on prior murine studies, our guinea pig data show that NE/IVT retains strong immunogenicity and protective efficacy in a transmission-relevant host. This is important because guinea pigs, unlike mice, support infection with unadapted human influenza isolates and efficient interhost transmission, partly due to a more human-like distribution of α2,6- linked sialic acid receptors in the upper respiratory tract (18, 20). Thus, our findings extend prior murine observations by showing that NE/IVT-adjuvanted mucosal vaccination can generate effective respiratory immunity in a model more suited for evaluating influenza transmission.

Our investigations demonstrate that, in the guinea pig model, IN triHA/NE/IVT vaccination induces systemic humoral responses comparable to IM triHA/Advx while providing a clear advantage in mucosal IgA induction at local and distal respiratory sites. Although IM triHA/Advx generated significantly elevated serum IgA titers, IN triHA/NE/IVT produced a more compartmentalized mucosal antibody profile, with detectable serum IgA in only 50% of animals. This discrepancy likely reflects the distinct mechanisms governing IgG and IgA transport into respiratory secretions. Mucosal IgG can be derived in part from circulating IgG through serum transudation and FcRn-mediated epithelial transport, linking mucosal IgG levels relatively closely to serum IgG titers (21, 22) In contrast, circulating IgA is predominantly monomeric and is not efficiently transported by FcRn, whereas secretory IgA is generated locally by mucosal plasma cells producing J-chain-containing polymeric IgA for pIgR-mediated transcytosis (9, 21, 22). Thus, high serum IgA does not predict robust mucosal IgA, as strong mucosal IgA more directly reflects local IgA induction (9, 23, 24). Consistent with this mechanism, IN triHA/NE/IVT vaccination induced the highest IgA titers in nasal washes, supporting effective antigen-specific IgA production at the respiratory port of entry. Respiratory secretory IgA is an important correlate of protection because it can neutralize, agglutinate, and entrap pathogens at mucosal surfaces, thereby limiting epithelial infection, viral shedding, and transmission while minimizing inflammatory effector functions associated with IgG-mediated complement or Fc receptor activation (9, 23–25). Although nasal wash neutralizing titers were modest, detectable local neutralizing activity in 50% of IN triHA/NE/IVT animals suggests that mucosal IgA contributes to local viral blockade. Together, these findings indicate that IN triHA/NE/IVT induces a distinct antibody response characterized by systemic humoral immunity together with targeted local IgA production, establishing first-line respiratory protection beyond passive entry of serum-derived antibodies.

An unexpected finding in this study was that IM triHA/AddaVax vaccination induced detectable antigen-specific IgG and IgA in mucosal samples, despite the absence of local antigen delivery. This response was not HA-specific, as a similar pattern was observed after IM vaccination with an unrelated antigen, trimeric SARS-CoV-2 spike protein, formulated with AddaVax. These results suggest that potent squalene-based IM adjuvantation may promote mucosal detection of antigen-specific antibodies in guinea pigs, potentially through serum-derived antibody transfer, vascular leakage, and/or migration of antibody-secreting cells to mucosal tissues. However, because vaccine-specific monomeric and dimeric/polymeric IgA cannot currently be distinguished in guinea pig mucosal samples, we cannot determine whether the detected IgA represents locally generated secretory IgA, serum-derived IgA, or both. This observation contrasts with mouse studies, in which comparable IM vaccination regimens generally induce minimal or undetectable respiratory mucosal antibodies (8, 19, 26). Thus, guinea pigs and possibly other larger animal models may be more permissive than mice to parenteral vaccine-driven mucosal antibody responses, suggesting that murine models may underestimate this contribution in other species.

Because most currently licensed influenza vaccines are administered intramuscularly, they primarily induce systemic humoral immunity but often generate limited immune responses in the upper respiratory tract. To address this limitation, IN boosting after IM priming—the so-called “prime-pull” strategy—has been proposed as a means to reinforce local mucosal immunity at the respiratory portal of viral entry (7, 8). Consistent with this concept, our data show that IN boosting with triHA/NE/IVT following QIV priming markedly enhanced both IgG and IgA responses in nasal washes, while maintaining serum IgG titers comparable to those induced by IM QIV vaccination alone. Notably, the IN-boosted group also exhibited higher microneutralization titers against Victoria/22, a strain included in the QIV prime but antigenically distinct from the triHA antigen (Michigan/15) used for IN boosting. This suggests that adjuvanted IN boosting may not only strengthen local antibody responses but also broaden or redirect pre-existing recall immunity toward antigenically related influenza strains. This mucosal enhancement was reflected in the challenge results. Only 25% and 50% of animals in the IM QIV; IN triHA/NE/IVT group shed infectious virus and viral RNA, respectively, and no transmission to co-housed naïve animals was detected. In contrast, all animals in the IM QIV-only group shed both infectious virus and viral RNA, and transmission occurred in one naïve contact. These findings indicate that IN boosting after systemic priming can improve upper respiratory viral control and reduce transmission beyond that achieved by IM vaccination alone. Although prior QIV priming may shape recall responses to heterologous mucosal antigens (27), the enhanced neutralizing activity against Victoria/22 despite boosting with antigenically distinct triHA suggests that this strategy may partially overcome limitations of conventional recall responses. Together, these results support triHA/NE/IVT as an effective mucosal booster after licensed IM influenza vaccination by enhancing local antibody responses, improving cross-reactive neutralization, and reducing viral shedding and onward transmission.

Interestingly, the mucosal antibody responses induced by IN triHA/NE/IVT vaccination were not restricted to the nasal cavity, the site of inoculation. Instead, NE/IVT strongly promoted antigen-specific IgA responses in nasal washes as well as at distal mucosal sites, including the oral and vaginal mucosa. This suggests that IN vaccination can engage a broader mucosal immune network and disseminate IgA-mediated immunity beyond the upper respiratory tract (9, 28, 29). This broad mucosal response may be particularly relevant because respiratory viruses are not always confined to the respiratory tract (30). Influenza infection can be associated with gastrointestinal symptoms, especially in children or during infection with novel influenza A viruses (31), while SARS-CoV-2 and adenoviruses can involve gastrointestinal (32), ocular (33), or other mucosal tissues. Therefore, distal mucosal IgA induced by nasal immunization may provide an additional layer of protection by limiting viral replication or subclinical infection at non-respiratory mucosal sites. Together, these findings suggest that NE/IVT-adjuvanted IN vaccination may strengthen immunity not only at the respiratory portal of entry but also across multiple mucosal compartments, thereby broadening protective coverage against respiratory viruses with extra-respiratory mucosal involvement.

The direct-contact transmission model provides strong evidence that adjuvanted mucosal vaccination can interrupt influenza spread. IN triHA/NE/IVT nearly eliminated infectious virus shedding in donor animals and completely prevented infection of co-housed sentinels, whereas unadjuvanted IN triHA failed to control replication or transmission. These findings indicate that IN antigen delivery alone is insufficient and that NE/IVT-mediated mucosal adjuvantation is critical for transmission-blocking immunity. The IN Michigan/15 group also showed strong protection, consistent with the robust mucosal and recall immunity induced by prior respiratory infection. However, because infection-induced immunity arises in the context of replicating virus, local inflammation, and broader immune priming, it should be viewed as a protective benchmark rather than an ideal strategy. Overall, the strongest protection was associated with conditions that enhanced respiratory mucosal immunity.

This study has several limitations. First, the group size was modest (n = 4), as is common in guinea pig transmission studies, which may have limited detection of subtle differences among vaccine regimens. Second, our analysis focused on H1N1 and a limited panel of viral strains, so protection against additional subtypes or more antigenically distant strains remains to be evaluated. Third, we did not directly assess T-cell responses, innate immune signatures, or mucosal B-cell and plasma cell populations, in part because guinea pig-specific immunological reagents remain limited. In addition, while the guinea pig model is highly valuable for influenza transmission studies, its respiratory tract anatomy and mucosal immune system are not identical to those of humans; therefore, extrapolation to human vaccine performance should be made with caution (15, 34). Finally, this study evaluated relatively short-term outcomes after vaccination and challenge. Future studies examining durability, broader strain coverage, longitudinal mucosal responses, cellular immunity, and cross-species comparisons will help define the generalizability of these findings and guide further development of NE/IVT as a modular mucosal vaccine platform.

Despite these limitations, our data provide strong proof-of-concept that NE/IVT-adjuvanted IN vaccination can induce potent systemic and mucosal immunity, broaden neutralizing activity when used as a booster after standard QIV, and markedly reduce or prevent influenza virus transmission in a relevant animal model. IN triHA/NE/IVT elicited systemic humoral responses comparable to parenteral vaccination while uniquely promoting robust local IgA responses at the respiratory portal of entry, nearly eliminating infectious viral shedding and preventing infection of co-housed naïve sentinels. Moreover, the ability of NE/IVT to function both as a homologous mucosal platform and as a heterologous IN booster highlights its potential value for next-generation influenza vaccines, including universal or supraseasonal strategies. Future studies should define the mechanisms underlying enhanced breadth, mucosal targeting, and transmission-blocking immunity, and further evaluate NE/IVT compatibility with diverse antigen designs, dosing regimens, and pre-existing immunity scenarios.

## Materials and methods

### Cells and Viruses

Madin-Darby Canine Kidney (MDCK) cells were cultured in Dulbecco’s Modified Eagle Medium (DMEM) supplemented with 10% fetal bovine serum (FBS), 100 IU/mL penicillin, and 100 μg/mL streptomycin.

Influenza virus stocks—comprising A/Netherlands/602/2009, A/Michigan/45/2015, and A/Victoria/4897/2022— were propagated in the allantoic cavity of 10-day-old embryonated chicken eggs at 37°C for 48 hours. Following harvest, the viruses were purified by ultracentrifugation through a 30% (w/v) sucrose cushion to ensure particle integrity and concentration. The resulting purified viral pellets were resuspended in phosphate-buffered saline (PBS) and stored at -80°C in single-use aliquots

### Animals

5-6 weeks old female Dunkin-Hartley guinea pigs (n=4/group) were obtained from Charles River Laboratories. Guinea pigs were transferred to BSL2 facilities and acclimated for 72 h before inclusion in the study. Ad libitum food and water was provided on a 12 h light/dark cycle.

### Immunization

Before immunization, guinea pigs were anesthetized by intraperitoneal injection of ketamine/xylazine at 30 mg/kg and 2 mg/kg, respectively. Animals received prime and boost immunizations 3 weeks apart. IN immunizations were administered in a total volume of 100 μL, with 50 μL delivered into each nare, whereas IM immunizations were administered in a total volume of 150 μL into the gluteal muscle of the left hind limb. A mock-vaccinated control group received IN PBS at both immunization time points and served as a negative control for antigen- and adjuvant-specific responses. The unadjuvanted IN triHA group received 30 μg of triHA protein alone at both prime and boost. To assess the effect of mucosal adjuvantation, a second group received 30 μg triHA formulated with 20% (w/v) nanoemulsion (NE) and 0.5 μg in vitro-transcribed defective interfering RNA (IVT DI RNA) by the IN route. For comparison with parenteral vaccination, a third group received 30 μg triHA formulated with 50% (v/v) AddaVax by the IM route. To assess whether mucosal adjuvanted triHA could boost pre-existing systemic vaccine immunity, guinea pigs were primed IM with QIV and boosted 3 weeks later with IN triHA/NE/IVT. A separate group received IM QIV at both prime and boost as a standard-of-care comparator. Each animal received 12 μg total HA from the 2023–2024 QIV formulation (3 μg per strain), which contained A/Victoria/4897/2022-like virus, A/Darwin/6/2021- like, B/Austria/1359417/2021-like, and B/Phuket/3073/2013-like viruses. To model infection-induced immunity, an additional group consisted of guinea pigs previously infected IN with seasonal H1N1 influenza virus and allowed to recover. In a separate cohort, guinea pigs were immunized IM with trimeric SARS-CoV-2 spike protein (triSpike), with or without AddaVax, to determine whether AddaVax-associated mucosal antibody responses reflected a broader adjuvant effect rather than HA-specific immunity. These animals received 30 μg triSpike in a total volume of 150 μL, were vaccinated twice at a 3-week interval, and serum and mucosal samples were collected 3 weeks after boost.

### Microneutralization assays

Microneutralization assays were performed as previously described (35). Briefly, MDCK cells were seeded at 1.4 × 10^4^ cells per well in 96-well plates and incubated overnight. RDE-treated serum, nasal wash, and bronchoalveolar lavage fluid samples were diluted 1:10, serially diluted two-fold in Opti-MEM, and mixed with influenza virus containing 100 TCID_50_. After 1 h incubation at room temperature, virus–sample mixtures were added to washed MDCK monolayers and incubated for 1 h at 37°C. The inoculum was removed, Opti-MEM containing TPCK-treated trypsin was added, and plates were incubated for 48 h at 37°C. Viral replication was assessed by hemagglutination using 0.5% chicken red blood cells, and neutralization titers were defined as the reciprocal of the highest dilution that completely inhibited hemagglutination.

### ELISA

Immunograde 96-well ELISA plates (Midsci) were coated with 50ng of recombinant trimeric HA protein in 50 μL PBS per well and incubated overnight at 4 °C. Plates were subsequently blocked with 200 μL of 5% non-fat dry milk in PBS for 1 hour at 37 °C. Samples collected from immunized guinea pigs were serially diluted in PBS containing 2% bovine serum albumin (BSA). Following removal of the blocking solution, diluted samples were added to the plates and incubated for 2 hours at 37 °C, followed by an overnight incubation at 4 °C. Plates were washed with PBST (PBS containing 0.05% Tween 20), and alkaline phosphatase–conjugated secondary antibodies diluted in PBS/2% BSA were added. The following secondary antibodies were used: HRP - donkey anti-guinea pig IgG (1:2,500, EMD Millipore: AP193P), HRP – rabbit anti-guinea pig IgM (1:2,500, Bioss: bs-0375R-HRP), and HRP -sheep anti-guinea pig IgA (1:2,500, ThermoFisher: SA5-10320). After a 1-hour incubation at 37 °C and wash, Plates were developed with TMB Turbo, stopped with 2N sulfuric acid, and absorbance was measured at 450 nm. Endpoint titers were calculated using a cutoff defined as the mean absorbance of the lowest dilution of naïve serum plus two standard deviations.

### Viral Challenge and Monitoring

Before viral challenge, guinea pigs were anesthetized as described above. Virus inocula were prepared in 300 μL PBS and administered intranasally, with 150 μL delivered to each nare. Body weight and rectal temperature were measured daily to monitor clinical responses after challenge. For transmission studies, inoculated donor animals were transferred to clean cages and co-housed with unimmunized naïve sentinels 8 h post-challenge for a 5-day observation period. To minimize cross-contamination, sentinel animals were handled before donors, gloves were changed between animals, and work surfaces were disinfected between cages. Nasal wash samples were collected at −1, 1, 3, and 5 DPI to assess nasopharyngeal viral loads over time. Under anesthesia, nasal washes were performed by instilling a total of 1 mL PBS into the nostrils and collecting the recovered wash fluid in a sterile Petri dish.

### Quantification of viral load and titers

Viral RNA was extracted from 150 μL nasal wash using the E.Z.N.A.® Viral RNA Kit (Omega Bio-tek) and quantified by Luna One-Step RT-qPCR (NEB) using 100 ng RNA per reaction. Viral copy numbers were calculated from a standard curve generated with 10-fold serial dilutions of an influenza A H1N1 nucleoprotein DNA template using nucleoprotein-specific primers (forward, 5′-GTTATGGCAGCATTCAGCGG-3′; reverse, 5′- GGTTCGTTGCCTTTTCGTCC-3′) and normalized to the starting nasal wash volume.

Infectious virus titers in nasal washes were determined by plaque assay as previously described (36). Briefly, clarified samples were serially diluted 10-fold in PBS, inoculated onto confluent MDCK monolayers, overlaid with agar-containing infection medium supplemented with TPCK-treated trypsin, and incubated for at least 48 h. Plates were then fixed, immunostained, and plaques were counted to determine infectious virus titers.

### Histopathologic analysis

At designated dpi, guinea pigs were euthanized and necropsied, and lung tissues were collected for histopathological analysis. Samples were fixed in 10% neutral buffered formalin, processed routinely, and embedded in paraffin. Paraffin-embedded tissues were sectioned at 6 µm using a microtome and mounted on glass slides. Sections were stained with hematoxylin and eosin (H&E) according to standard protocols. Histopathological evaluation was conducted by a pathologist in a double-blinded manner. Detailed scoring criteria are provided below.

**Table 1.**

| Parameter | Histopathologic Score |  |  |  |
| --- | --- | --- | --- | --- |
|  | 0 | 1 | 2 | 3 |
| Peribronchiolar inflammation | None | Mild, Loosely formed cuffs of inflammatory cells | Moderate, Well-formed cuffs of inflammatory cells | Prominent thick well-formed cuffs of inflammatory cells |
| Perivascular inflammation | None | Mild, Loosely formed cuffs of inflammatory cells | Moderate, Well-formed cuffs of inflammatory cells | Prominent thick well-formed cuffs of inflammatory cells |
| Bronchiole lumen inflammation | None | Low | Medium | High |
| Alveolar space inflammation | None | Mild | Moderate | Severe |
| Broncho-interstitial pneumonia | None | Mild | Moderate | Severe |

## Data availability

All data files are available for immediate release upon request from the corresponding authors.

## Acknowledgements

Research in the M.S. laboratory is funded by NIH/NIAID grants R21AI180874, R21AI176069, R01AI160706, and partly funded by CRIPT (Center for Research on Influenza Pathogenesis and Transmission), a NIH NIAID-funded Center of Excellence for Influenza Research and Response (CEIRR, contract number 75N93021C00014) and partly by CIVICs (NIAID contract # 75N93019C00051). V.Y. receives support through training grant T32AI007647 (Program Director Domenico Tortorella and co-director Viviana Simon). Research in the P.T.W. laboratory is funded by NIH/NIAID grants R21AI180874, R21AI176069, R01AI160706. We would like to thank Gowthamee Thangavel for her helpful guidance during the initial setup of the guinea pig experiments.

## Author contributions

V.Y., S.C.P., P.T.W. and M.S. conceived and designed the study. V.Y. and S.C.P. performed the experiments and analyzed the data. V.Y., S.C.P., P.T.W. and M.S. wrote the manuscript. V.Y., S.C.P., G.L., M.J.W., J.N.D., and C.C. performed experiments. J.L. and F.E. generated critical reagents for this study. P.T.W. and M.S. secured funding, provided overall project direction, and edited the manuscript with input from all authors. V.Y., S.C.P., M.W., P.T.W and M.S. reviewed and edited the manuscript for clarity and accuracy. All authors have read and approved the manuscript.

---

**Supplement Fig 1.**
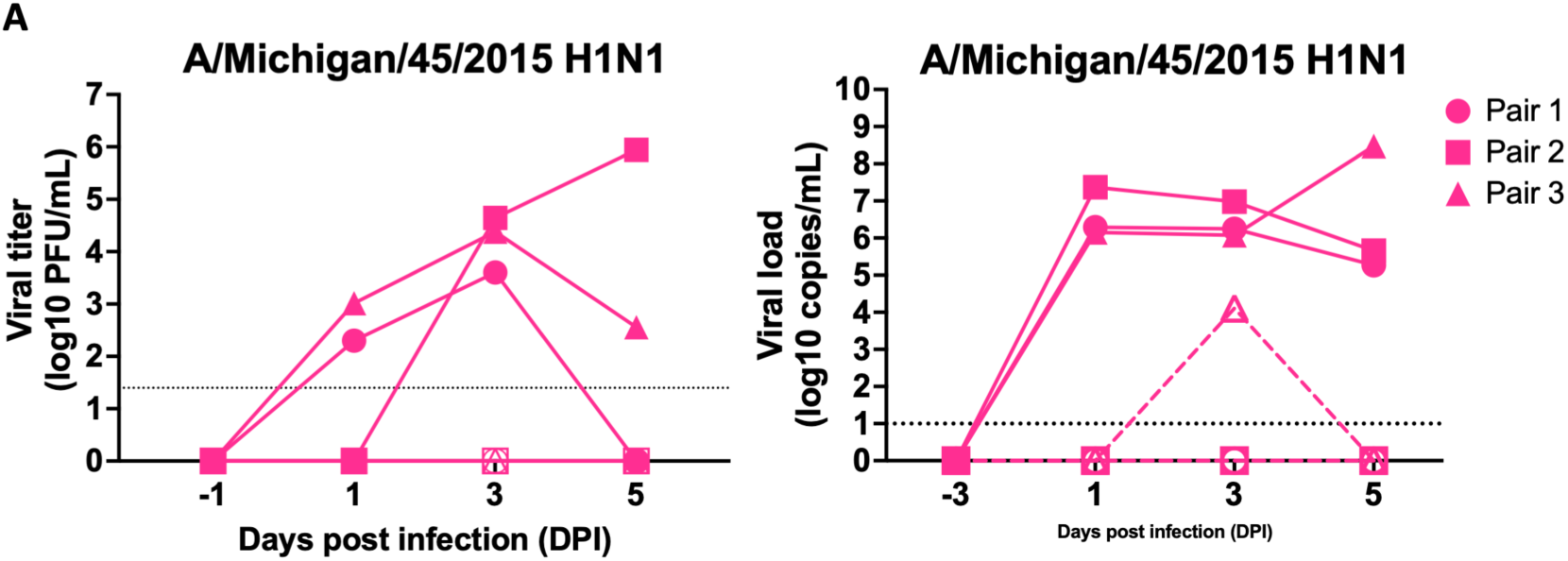
Limited transmission of A/Michigan/45/2015 in naïve guinea pig pairs. Naïve donor guinea pigs were intranasally inoculated with A/Michigan/45/2015 (H1N1) and co-housed with naïve sentinel animals 8 h later to assess direct-contact transmission. Nasal wash samples were collected at the indicated time points and analyzed for (left) infectious virus by plaque assay and (right) viral RNA by qPCR. Filled symbols with solid lines represent donor animals, and open symbols with dashed lines represent sentinel animals. Each symbol shape denotes an individual donor–sentinel pair. Dashed horizontal lines indicate the limit of detection of each assay. *, P < 0.05; **, P < 0.01; ***, P < 0.001; ****, P < 0.0001.

